# Pharmacokinetics of Extended-release Buprenorphine (Ethiqa) in Female Yorkshire Swine (*Sus scrofa domestica*)

**DOI:** 10.1101/2023.10.04.560917

**Authors:** Lynlee M. Stevey-Rindenow, Marissa Saenz, Vivian La, Craig Franklin, Ari Aycock-Williams, Patrick T. Fueger

## Abstract

Despite the use of swine as a large animal translational surgical model, precise dosing regimens for commonly used analgesics such as buprenorphine, are currently lacking in this species. A newly available extended-release formulation of buprenorphine (XRB, Ethiqa) is FDA-indexed and approved for use in mice and rats; however, no studies have examined the efficacy and pharmacokinetic parameters of XRB in swine. The goal of this study was to determine the pharmacokinetics of the newly available XRB in swine. We hypothesized that after a single subcutaneous administration of XRB in adult swine, buprenorphine plasma concentrations would be at or above the therapeutic threshold of 0.1 ng/mL and that local injection side effects would be minimal. XRB was administered once, subcutaneously to two separate cohorts of adult female Yorkshire swine at low and high doses (0.2 and 0.4 mg/kg, respectively; n = 3 and 2). Blood was collected from an indwelling jugular catheter prior to and after XRB administration (13 total time points). Individual animal data indicated all animals reached therapeutic buprenorphine plasma concentrations by 8 h post administration. Average plasma buprenorphine levels for both the low- and high-dose cohorts reached therapeutic concentrations starting at 1.5 h after XRB administration and were maintained above therapeutic concentrations throughout the 96-h study period. In the low-dose cohort, the average half-life was 212.6 ± 107.1 h, whereas the half-lives in the high-dose cohort was 63.8 and 48.9 h. As expected, histology of XRB subcutaneous sites revealed mild injection site reactions characterized by granulomatous inflammation with intralesional cholesterol cleft formation. These results support our hypothesis and indicate that all animals maintained therapeutic plasma buprenorphine levels beginning at 8 h and maintaining past 96 h. Thus, XRB at either dose provide therapeutic levels of plasma buprenorphine and therefore its use should be further explored in swine.

## INTRODUCTION

Swine are the most widely used large animal translational model in biomedical research.^43^ The anatomical and physiological similarities between swine and humans make them ideal for pharmaceutical and toxicology studies as well as surgical training. Although there have been significant refinements in the humane handling, anesthesia, and recovery monitoring of swine, precise dosing regimens for commonly used analgesics are currently lacking.^43,44,45,46^ To provide adequate animal welfare and pain management, the dose, frequency, and concentration of an analgesic must be appropriate.^56^ As a USDA-covered species, swine should be provided a safe, reliable, and efficacious pain management that is also largely free from adverse effects.

Buprenorphine is one of the most widely used opioid analgesics administered to both laboratory rodents and companion animals.^38^ As a semi-synthetic partial μ-opioid agonist, δ- and κ-opioid antagonist with action in the brain, spinal cord, and peripheral tissues, buprenorphine suppresses both pain transmission and modulation.^2,50^ Buprenorphine is highly lipophilic and exhibits a ceiling effect with slow binding of the μ-opioid receptor providing a long duration of action for analgesia – important qualities in effective and safe analgesia.^10,51,52^ Although dose-dependent respiratory depression and sedation are possible, many studies have established buprenorphine to be safe and effective in common laboratory animal species including mice, rats, and non-human primates.^1,4,12,14,16,27,33,38^

Despite the plethora of information regarding buprenorphine use in rodents, there is little information on adequate buprenorphine use in swine. The reported dose for buprenorphine HCl in swine is wide and ranges from 0.01-0.1 mg/kg with a dosing frequency of twice or three times daily.^8,13,17,18,20,30,37,40,45,46,50,55^ The long-lasting buprenorphine formulations, formerly sustained-release buprenorphine (SRB; Zoopharm, Fort Collins, CO) and the newly available extended-release buprenorphine (XRB, Ethiqa; Fidelis Pharmaceuticals, North Brunswick, NJ), are superior to the conventional buprenorphine HCl formulations because their long-lasting formulations avoid repeat administrations – an important refinement for minimizing animal distress and pain. However, SRB pharmacokinetics in swine has only been described in two studies with reported dose ranges from 0.12-0.24 mg/kg subcutaneous, once daily and therapeutic plasma concentrations of 0.1 ng/mL or more for at least 96 h.^17,47^ To date, no studies have examined the pharmacokinetic profile of XRB in swine.

The goal of this study was to characterize the pharmacokinetics of the newly available XRB, (Ethiqa) in swine. We hypothesized that after a single subcutaneous administration of Ethiqa in adult Yorkshire swine, buprenorphine plasma concentrations would be at or above the therapeutic threshold of 0.1 ng/mL. Furthermore, we hypothesized that subcutaneous Ethiqa administration would result in minimal, serious local side effects at the site of injection.

## MATERIALS AND METHODS

All animal handling and procedures were approved by the AAALAC International accredited University of Southern California Institutional Animal Care and Use Committee (IACUC). All animals were housed in accordance with the *Guide for the Care and Use of Laboratory Animals*, Public Health Service Policy on the Humane Care and Use of Laboratory Animals, and the Animal Welfare Act and Regulations.^49,56^

### Animals

Six adult female Yorkshire domestic (*Sus scrofa domestica*) swine approximately 3-months-old, weighing on average 30-35 kg were obtained from Premier BioSource (formerly S&S Farms, CA), a closed herd and Class A USDA-licensed facility. Although six animals were initially enrolled in the study, only five were used due to kinks in the catheters that prohibited blood collection of one animal. Animals were maintained under strict barrier conditions, monitored for pathogens quarterly, and regularly screened for parasites. According to the vendor provided health monitoring quarterly reports, all animals were free from major swine bacterial, viral, and parasitic pathogens including but not limited to: Porcine epidemic diarrhea virus, Porcine reproductive and respiratory syndrome virus, *Actinobacillus pleuropneumonia*, transmissible gastroenteritis virus, Mycoplasma, Influenza A, Parvovirus, Leptospirosis, Brucellosis, and Pseudorabies.

### Facility

Animals were fed pelleted feed (Laboratory Porcine Diet Grower 5084, Lab Diet) twice daily, and food enrichments such as fruit and vegetables every other day. Environmental enrichment included scratch pads, manipulatable malleable toys, and hanging foraging balls and were available on a rotating basis. Water was provided *ad libitum* from an in-kennel automatic watering system. Animal room was maintained at 70-73°F with 30-70% relative humidity with 100% recirculated air at 10-15 changes hourly. Fluorescent lighting was provided on a 12:12-h light: dark cycle. Animals were housed in stainless steel bar type open top kennels with raised floors. Prior to enrollment in the study, animals were deemed healthy as determined by veterinary physical examination, comprehensive complete blood count, chemistry, and fecal examination. Animals were monitored daily throughout the study by the veterinarian and monitored after hours via a Wi-Fi enabled live video monitoring system placed inside each animal’s home kennel.

### Acclimation

To facilitate low stress sedation, recovery period and subsequent serial blood collections, the animals were acclimated to close contact with research staff via gentle touch and petting of the neck and ear region, high value food rewards, and use of a Panepinto sling.^35,36^ Acclimation to a Panepinto sling was accomplished through a previously described technique with multiple short (5-30 min) daily training sessions and positive reinforcement for a minimum of 7-10 days before the start of study.^54^

### Experimental Design

All animals enrolled in this study were part of an IACUC approved protocol with the Department of Surgery at Keck Medicine of USC. Animals were co-housed in pairs prior to the start of the study and during acclimation period. After catheterization and for the remainder of the study, animals were then singly housed in adjacent kennels while maintaining the ability to interact with each other through sniffing, vocalization, and sight through kennel bars. Animals were grouped in pairs and each animal was randomly assigned to either a low dose (0.2 mg/kg, SC, once) or high dose (0.4 mg/kg, SC, once) administration of XRB (Ethiqa). Animals assigned the low dose group received a single injection of XRB (Ethiqa) in the subcutaneous space behind one ear, whereas an equal volume of normal saline (Hospira, Inc., Lake Forest, IL USA) was administered in the subcutaneous space of the contralateral ear as a control. Due to the large, calculated volume of the high dose, animals assigned to the high dose group received an evenly split dose (mL) administered in the subcutaneous space behind both the left and right ears. Animals were fasted beginning the night before catheterization, and catheters were placed under sedation (described below). After placement of the catheter, a time 0 blood sample was collected for baseline pharmacokinetic analysis and animals were administered either the low or high dose of XRB (Ethiqa). Blood was subsequently collected from animals at 0.25, 0.5. 1, 1.5, 2, 4, 8, 12, 24, 48, 72, and 96 h post-administration. After completion of the study, animals were used in a separate terminal surgical training program procedure, as mentioned above. Post-mortem full thickness skin and subcutis tissue samples were collected at drug and saline control injection sites and placed in 10% neutral buffered formalin until paraffin embedding, slide preparation and staining with hematoxylin and eosin (slides prepared by IDEXX BioAnalytics, Columbia, MO).

### Catheters Placement

Animals were fasted a minimum of 12 h in preparation for sedation to perform jugular catheter placement and subsequent drug administration. Animals were sedated with a single intramuscular injection of Telazol (Zoetis Inc., Kalamazoo, MI, USA) at 50 mg/mL, Tiletamine HCl at 50 mg/mL, and Zolazepam HCl reconstituted with Xylazine (100 mg/mL) to make a final concentration of 100 mg/ml at a dose administration of 2.2-4.4 mg/kg body weight. Animals were maintained on inhaled isoflurane mixed with oxygen and delivered by precision vaporizer via face mask as needed to maintain anesthesia. A single triple lumen indwelling jugular catheter (7Fr x 30cm, J1045B; JorVet; Jorgensen Laboratories, Inc., Loveland CO, USA) was placed percutaneously and aseptically using previously described Seldinger technique, ultrasound guidance, and/or anatomical triangulation techniques in the right external jugular vein.^6,15,42^ At the time of jugular catheter placement, the luminal volume of the catheter was measured to be approximately 1 mL. Once placed, the jugular catheter was sutured to the skin and further secured with a sterile Tegaderm pad (Tegaderm Foam Dressing #90610, 3M Health Care, St. Paul, MN, USA). Cast padding, stockinette/stretch tubular elastic dressing retention net, Size 11 (Curad®, Medline, Mundelein, IL, USA), and Elastikon (Johnson & Johnson Consumer Companies, Inc. Skillman, NJ, USA) was used to make a comfortable ‘jacket’ around the upper torso and neck. Before and after all timed blood draws, catheters were flushed and locked with heparinized saline (100 U/mL; Becton, Dickinson and Company, Franklin Lakes, NJ, USA) to maintain patency. Although not used during this study, an additional catheter was placed in either the left or right marginal auricular vein as a backup to the jugular catheter to ensure venous access.

### Behavioral Scoring

Given opioids are analgesics can synergistically act with other drugs to induce either hyperactivity and/or extreme sedation in other species, we monitored behavioral parameters for sedation and distress. A sedation and distress scoring guide (Figure 1), adapted from *Santos et al*. was used to determine the degree of sedation and distress in each animal after administration of low or high dose XRB (Ethiqa).^9,41^ Scoring was assessed cage side, at time 0 and 0.25, 0.5. 1, 1.5, 2, 4, 8, 12, 24, 48, 72, and 96 h post-administration of XRB (Ethiqa). All animals were monitored multiple times daily by a veterinarian and remained healthy throughout the study. A second, separate veterinarian blinded to the treatment groups evaluated the overall welfare of animals approximately 24-48 h post drug administration. Veterinary exams did not reveal any serious adverse effects.

**Figure 1.**
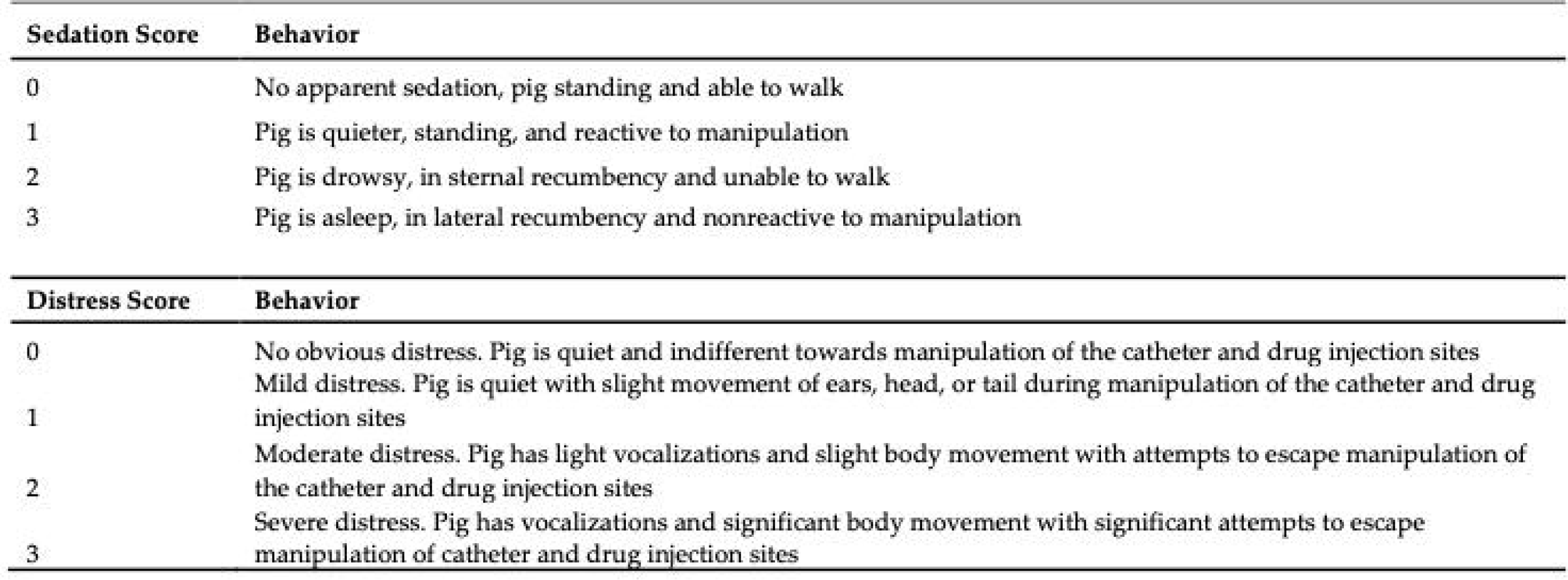
Sedation and distress scoring system clinically evaluated in swine up-to 96 hours after XRB (Ethiqa) administration (adapted from Santos et al.^41^).

### Drug Administration and Blood Collection

At the conclusion of jugular catheter placement and securement, XRB (Ethiqa) low dose (0.2 mg/kg) or high dose (0.4 mg/kg) was administered. Animals were allowed to recover from sedation in a Panepinto sling for approximately 1 to 2 h post drug administration. Animals were then slowly transitioned to either a rolling padded kennel for additional safe recovery or their home kennel depending on level of recovery. Approximately, 1 mL of the heparinized saline lock (previously measured) was collected via the jugular catheter injection port using an 18G butterfly catheter (Surflo, Terumo Corporation, Hatagaya, Shibuya-ku, Tokyo, Japan) attached to a 3 mL syringe to clear the catheter prior to blood sample collection. A new 18G butterfly catheter attached to a new 3 mL syringe was then used to draw the experimental 1-2 mL blood sample and was immediately transferred to EDTA tubes (Greiner Bio-One, Monroe, NC, USA) at times 0.25, 0.5. 1, 1.5, 2, 4, 8, 12, 24, 48, 72, and 96 h post-administration of XRB (Ethiqa). Blood samples were gently and repeatedly rolled in EDTA tubes and placed at 4□ immediately after collection until centrifugation at 3,800 x *g* for 10 min at 4□. Plasma was then collected and subsequently stored at -80□ until analysis approximately 1 month post collection. Plasma was then shipped overnight on dry ice to the McWhorter School of Pharmacy, Pharmaceutical Sciences Research Institute in Samford University, Birmingham Alabama, for measurement of plasma buprenorphine concentrations using liquid chromatography tandem mass spectrometry (HPLC MS/MS). A presumed therapeutic plasma concentration of 0.1 ng/mL was used in this study and based on the limited injectable buprenorphine pharmacokinetic data in both swine and extrapolations from human and non-human primate literature.^16^

### Pharmacokinetic Analysis

Buprenorphine standard spiking solutions were prepared in 50:50 DI water: acetonitrile to give concentrations in plasma ranging from 0.2-200 ng/mL. The buprenorphine plasma samples and standards (100 μL) were fortified with internal standard (50 ng/mL terfenadine). Acetonitrile (1 mL) was added to precipitate the plasma proteins, and the mixture was vortexed and centrifuged. The organic layer was transferred to a clean test tube and evaporated to dryness under nitrogen in a water bath set at 50°C. The samples were reconstituted in dilution solvent and analyzed by HPLC MS/MS. Matrix matched standards and QC samples were prepared using blank control plasma. Chromatographic separation of buprenorphine and the internal standard from the extracted plasma matrix was achieved using a Shimadzu HPLC system consisting of two Shimadzu LC20-AD pumps, and SIL20-AC HT autosampler, and a DGU-20A3-3channel in-line degasser and Controller (Columbia, MD) with a 100 mm x 2 mm Luna C18 reverse phase column (Phenomenex, Torrance CA) at ambient temperature. The mobile phase consisted of 5 mM ammonium acetate and acetonitrile, each fortified with 0.1% formic acid. The compounds were analyzed using a gradient elution profile in which mobile phase B was held at 30% for 1 minute, then increased to 80% over 4 min, held at 80% for 0.5 minutes, then returned to 30% and equilibrated for 2.5 min. Mass detection was accomplished with an Applied Biosystems (Foster City, CA) 4000 QTRAP triple quadrupole ion trap mass spectrometer equipped with an electrospray ionization source operated at a potential of 5 kV at 450°C operating in the MRM mode. Data were collected and processed using Analyst 1.6.2 (Applied Biosystems, Foster City, CA). The following mass transitions of the compounds were monitored: buprenorphine, m/z 468.4 – 396.1; and terfenadine, 472.4 – 436.2. Data were analyzed using a Noncompartmental Analysis.

## RESULTS

### Pharmacokinetics

The pharmacokinetic data for animals treated with low- (0.2 mg/kg) or high- (0.4 mg/kg) dose XRB (Ethiqa) are reported in Figure 2 and Table 1. Regardless of dose, buprenorphine was undetectable in all samples from the 15 min post drug administration time point. Average plasma buprenorphine levels for both the low- and high-dose cohorts reached therapeutic concentrations of 0.1 ng/mL starting at 1.5 h after XRB (Ethiqa) administration and were maintained above therapeutic concentrations throughout the 96-h study period (Figure 2C). In the low-dose cohort, the average half-life was 212.6 ± 107.1 h, whereas the half-lives in the high-dose cohort were 63.8 and 48.9 h (Table 1). The earliest time point in which plasma concentration reached the therapeutic threshold was at 90 min in animal B (high dose, Figure 2B) and the latest time point was at 8 h in animal F (low dose, Figure 2A). Both animals assigned to high dose XRB (Ethiqa) (Figure 2B) reached therapeutic concentrations earlier than animals in low dose group - animal B at 1.5 h and animal C at 2 h (Figure 2B). In contrast, low dose animals reached the therapeutic concentration at 4 h for animal A, 1.5 h for animal E, and 8 h for animal F (Figure 2A). XRB (Ethiqa) was present in the plasma at the therapeutic concentrations of 0.1 ng/mL or above at time points beginning at 8 h and maintaining throughout the 96-h time point for all animals in both groups. Although the time to peak concentration varied between individual animals within each dose group, the peak buprenorphine plasma concentration for low-dose group was consistent with an average of 0.36 ± 0.03 ng/mL (Table 1). The peak buprenorphine plasma concentration for the high-dose group varied with animal B at 1.14 ng/mL and animal C at 0.73 ng/mL. With a mean AUC of 23.8 ± 1.9 and 48.1 h x ng/mL in the low- and high-dose XRB (Ethiqa) groups, respectively, the high-dose XRB (Ethiqa) group was doubled that of the low-dose group. In addition, the clearance of the drug varied between animals in both the low- and high-dose groups (Table 1).

**Table 1.** Pharmacokinetic parameters of low- and high-dose XRB (Ethiqa)-treated pigs were calculated from a non-compartmental pharmacokinetic analysis^1^ (NCA) of averaged buprenorphine concentrations. AUC _0-last_ = area under the concentration-time curve. Data for low-dose XRB (Ethiqa) (n=3) are presented as mean ± SEM and data for high-dose XRB (Ethiqa) (n=2) are presented as a mean.

|  | Low-dose<br>(0.2 mg/kg, SC) |  |  |  | High-dose<br>(0.4 mg/kg, SC) |  |  |
| --- | --- | --- | --- | --- | --- | --- | --- |
| Individual Pig | A | E | F | Avg. ± SE | B | C | Avg. |
| Half-life (h) | 169.2 | 415.9 | 52.6 | 212.6 ± 107.1 | 63.8 | 48.9 | 56.4 |
| Time to peak concentration (h) | 24 | 2 | 48 | 25 ± 13 | 4 | 48 | 26 |
| Peak concentration (ng/mL) | 0.35 | 0.42 | 0.32 | 0.36 ± 0.03 | 1.14 | 0.73 | 0.9 |
| AUC<sub>0-last</sub> (h × ng/mL) | 25.4 | 26.1 | 20.0 | 23.8 ± 1.9 | 47.3 | 48.8 | 48.1 |
| Clearance (mL / h / kg) | 2381 | 769 | 6077 | 3076 ± 1517 | 5213 | 5,338,826 | 2,672,019 |

**Figure 2.**
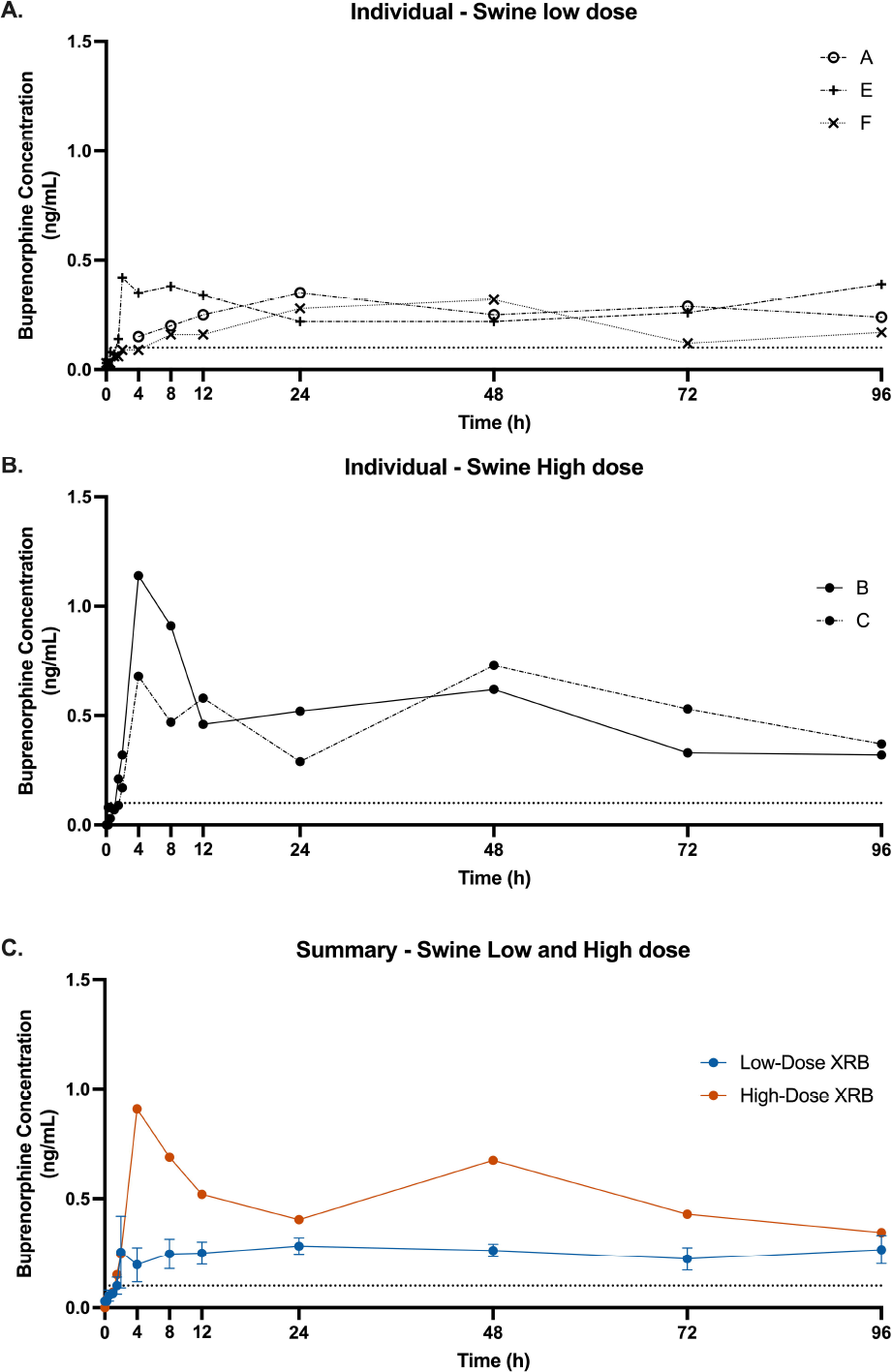
Swine treated with low-dose (0.2 mg/kg, SC) or high-dose XRB (Ethiqa) (0.4 mg/kg, SC) had circulating buprenorphine levels that were above the therapeutic effective threshold of 0.1 ng/mL (dashed line). Individual animal low-dose (A) and high-dose (B) circulating buprenorphine levels up to 96 hours post-administration. Averaged circulating buprenorphine levels (C) of low-dose (blue line) or high-dose (red line) XRB (Ethiqa)-treated pigs. Data for low-dose XRB (Ethiqa) (n=3) are presented as mean ± SEM and data for high-dose XRB (Ethiqa) (n=2) are presented as a mean.

### Sedation and Distress Scores

Although distress and sedation ethograms for adult swine have not been established, a sedation scoring system has been proposed in piglets by Di Giminiani *et al*.^9^ For this study, we used an adapted form of Santos *et al*.^41^ sedation scoring system to determine the degree of distress and sedation in these catheterized adult swine (Figure 1). As expected, sedation scores (from 0-3, with 0 being no apparent sedation and 3 being asleep in lateral recumbency) peaked at the time of administration for both low- and high-dose XRB (Ethiqa) animals (Table 2). The last animal to reach an un-sedated state (score of 0) was animal B at 24 h (high dose, Table 2). Sedation scores for both low and high dose groups did not correlate with Tmax or Cmax. The data indicate that distress scores (from 0 to 3, with 0 being no apparent distress and 3 being severe distress with vocalization and body movement) peaks variably between animals and doses. Low-dose animals A and E have peak distress scores ranging from 1.0 to 1.5 at 0.5 h – 2 h, while animal F did not appear distressed at any point in the study. Among the high-dose animals, animal B had a distress score of 3 at 1.5 h – 2 h and did not appear normal until 24 h after XRB (Ethiqa) administration and animal C had peak distress scores at 1 h – 1.5 h (Table 2). Interestingly, animal B sedation and distress scores both appear back to normal (score of 0) by 24 h post high-dose XRB (Ethiqa) administration (Table 2).

**Table 2.**
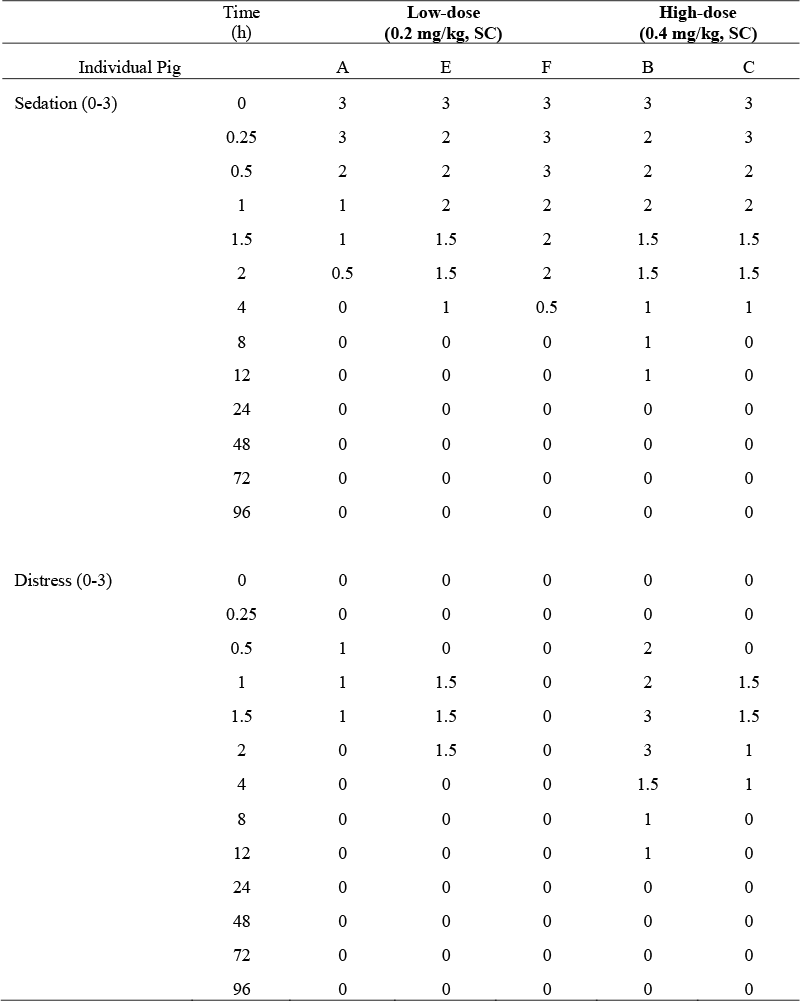
Sedation and distress scores in individual pigs up to 96-hours after low- or high-dose XRB (Ethiqa) administration.

### Histopathology

Gross skin lesions were not apparent during biopsy collection in either low- or high-dose XRB (Ethiqa) subcutaneous injected sites. Saline injected sites revealed no abnormal lesions grossly or histologically. Both low- and high-dose XRB (Ethiqa) biopsies revealed a thick white-to-cream colored tissue reaction that extended down the subcutis; the high-dose XRB (Ethiqa) tissue reactions extended past the subcutis and into the musculature. Histologic evaluation of XRB (Ethiqa) injection sites revealed similar lesions in all pigs regardless of dose (Figure 3, Table 3). The deep subcutaneous fat was characterized by varying degrees of granulomatous inflammation with patterns varying from dense macrophage infiltrates with frequent giant cell formation and numerous intralesional cholesterol clefts to large vacuoles lined by flattened epithelium and often rimmed by primarily macrophage infiltrates with varying numbers of accompanying neutrophilic infiltrates. Varying degrees of fibrosis were also detected amidst these vacuoles. The vacuolated pattern was usually at the periphery of the denser inflammation and inflammation and vacuolation were often branching into adjacent superficial fatty tissue and deep musculature. Within the dense inflammation single to multiple cores of mineralized necrotic debris rimmed by degenerating cells including neutrophils were observed in some swine, as were pockets of hemorrhage. A representative slide was evaluated under polarized light and no birefringent material (e.g., plant material) was detected.

**Table 3.** Characteristic lesions found in H&E-stained skin biopsies from low- and high-dose XRB (Ethiqa) treated pigs. Data are presented as a proportion with a total of n=3 for low-dose XRB (Ethiqa)-treated pigs and a total of n=2 for high-dose XRB (Ethiqa)- treated pigs (*hemorrhage from pig B was extensive).

|  | Low-dose<br>(0.2 mg/kg, SC) | High-dose<br>(0.4 mg/kg, SC) |
| --- | --- | --- |
| Granulomatous inflammation | 3/3 | 2/2 |
| Cholesterol clefts / vacuoles | 3/3 | 2/2 |
| Fibrosis | 3/3 | 2/2 |
| Neutrophil infiltration | 3/3 | 2/2 |
| Hemorrhage | 2/3 | *2/2 |
| Mineralization of necrotic debris | 2/3 | 1/2 |

**Figure 3.**
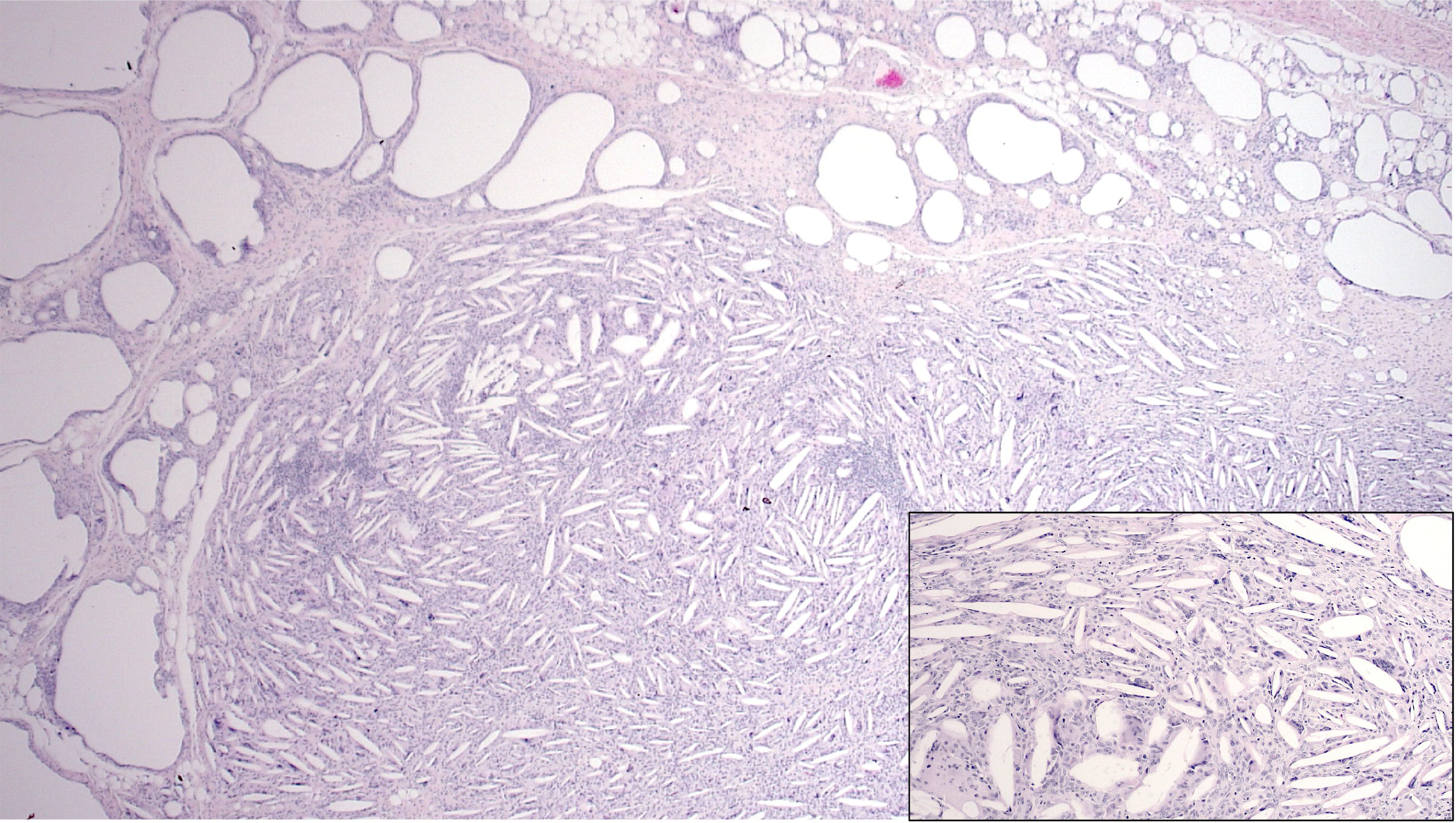
Representative lesion seen in deep subcutaneous fat of pigs injected with XRB (Ethiqa) regardless of dose showing dense granulomatous inflammation with numerous intralesional cholesterol clefts bordered by large vacuoles lined by flattened epithelium and often rimmed by macrophage infiltrates. Inset shows a higher power of dense granulomatous infiltrates with cholesterol clefts and giant cell formation.

## DISCUSSION

We characterized the pharmacokinetics of the injectable suspension of extended-release buprenorphine, XRB (Ethiqa), in adult female Yorkshire swine. Furthermore, we described animal sedation and distress after XRB (Ethiqa) administration and assessed injection site reactions. This study is the first to reveal that XRB (Ethiqa) reaches the therapeutic buprenorphine plasma concentration of 0.1 ng/mL in adult swine at a low (0.2 mg/kg) and high (0.4 mg/kg) dose. All animals reached the therapeutic plasma threshold by 8 h and maintained it to the end of the study (96 h). Animals had minimal lesions at the XRB (Ethiqa) subcutaneous injection site and a transient change in sedation and distress scores that returned to normal by 24 h after administration. Together these results suggest that the extra-label use of XRB (Ethiqa) in adult swine provides safe and sustained analgesia.

XRB (Ethiqa) is currently the only pharmaceutical grade FDA-indexed extended-release buprenorphine formulation that is approved for use in mice and rats.^29,57^ In multiple strains of mice and rats, pharmacokinetic studies revealed a single injection of XRB (Ethiqa) provides therapeutic plasma concentrations for 48-72 h.^1,4,21,22,23,26,31,32,40^ Additionally, XRB (Ethiqa) has a superior safety profile and long-lasting analgesia effects of up-to 72 h in mice and rats after a single subcutaneous injection.^3,7,16,19,24,40,48^ Our results indicate in swine XRB (Ethiqa) provides longer therapeutic buprenorphine concentrations, consistent with what is observed in other non-rodent species and swine breeds. In the cynomolgus macaque (*Macaca fascicularis*) and the common marmoset (*Callithrix jacchus*), therapeutic buprenorphine plasma levels lasted as long as 96 h after XRB administration.^11,25^ Evaluation of another non-FDA approved long-lasting buprenorphine, SRB, in diabetic Yucatan miniature swine revealed therapeutic plasma levels that surpassed 96 h.^17^ Furthermore, our study revealed XRB (Ethiqa) injection sites sustained minimal changes including granulomatous inflammation and cholesterol clefts – consistent with what has been reported with injection of XRB (Ethiqa) in other species and likely due to the lipid-bound in a medium chain fatty acid triglyceride (MCT) oil.^11,16,22,23,25^ Therefore, XRB (Ethiqa) extra-label use in swine increases our options for long-lasting and safe analgesia in swine.

Unfortunately, other currently available opioid-based pain management options in swine provide a highly variable and unreliable dose range with fluctuations in therapeutic plasma concentrations and analgesic effectiveness.^18,28,34,45,47,53^ Although fentanyl patches are used, there is concern with inaccurate dosage associated with patch location, variability among species and breeds, analgesic effectiveness, and potential adverse effects.^18,28,34,45,53^ Transdermal forms of buprenorphine have been evaluated in only a few studies. One study indicated 30 μg/h patches in Yucatan minipigs had peak plasma buprenorphine concentrations between 0.37-0.75ng/mL at 12-24 h after application.^47^ Other studies found that two different transdermal patch doses (35 and 70 μg/h) failed to produce reliable serum buprenorphine concentrations; thus, transdermal patches are generally not recommended for use in swine^28,34^ Furthermore, although extra-label use of long-lasting injectable SRB in swine has been shown to reach therapeutic plasma concentrations, there are reservations with its safety, purity, and efficacy between batches since it has not undergone FDA approval.^56^ Studies comparing local injection reactions of SRB and XRB (Ethiqa) in multiple species revealed XRB (Ethiqa) had less inflammatory changes.^11,23,25^ These findings encourage further optimization of long-lasting buprenorphine formulation for swine and support the extra-label use of the FDA-indexed XRB (Ethiqa) in this species.

Our primary goal was to characterize the pharmacokinetic parameter of XRB (Ethiqa) in adult female Yorkshire swine; however, we acknowledge a few limitations to our study. The therapeutic buprenorphine plasma concentration of at least 0.1 ng/mL used in this study has not been validated in swine and is extrapolated from other species.^5,10,16,47,52^ However, it is well known that pharmacokinetic studies cannot provide the full physiologic or biological effects on the animal. Studies have recommended not making estimations of adequate analgesia based on the level of drug in plasma until a standardized, validated, and humane methodology using a pain model or analgesiometric test is assessed.^39,53^ Furthermore, although we did not assess XRB (Ethiqa) efficacy in a painful swine model but we were able to clinically assess potential adverse effects using novel sedation and distress ethograms adapted from an adult swine sedation ethogram.^9,41^ Overall, administration of postoperative analgesics should be based on multiple clinical factors including reaction to incisional pain, abnormal posture, reluctance to lie down, or appetite related changes.^44^

This study is the first to establish that a single subcutaneous administration of the newly available XRB (Ethiqa) in swine provides buprenorphine plasma concentrations above the therapeutic threshold of 0.1 ng/mL beginning at 8 h and maintaining past 96 h. Our results are consistent with our hypotheses and suggest that extra-label use of XRB (Ethiqa) does not result in serious adverse injection reactions or changes in clinical signs of sedation and distress. These results support the use of multimodal analgesics to ensure adequate pain management and increases our options of long-lasting analgesics in swine within the laboratory animal medicine field.

## Abbreviations and Acronyms

XRB: extended-release buprenorphine
SRB: sustained-release buprenorphine.

## ACKNOWLEDGMENTS

The authors would like to thank Dr. Adrian Dobrowolsky, Mrs. Angela Martinez, and Ms. Tiffany Barajas from the Department of Surgery at Keck Medicine of USC for animals enrolled in the study; Dr. Gregory Gorman at the McWhorter School of Pharmacy, Pharmaceutical Sciences Research Institute at Samford University for pharmacokinetics analysis; and Emi Katayama LATg and the ‘HMR Crew’ for their exceptional animal care of A, B, C, D, E, F. MS was supported by a National Cancer Institute Cancer Metabolism Training Program Postdoctoral Fellowship (T32CA221709). This work was supported, in part, by intramural funding from the Department of Animal Resources at the University of Southern California and the Beckman Research Institute, City of Hope. Fidelis Animal Health Inc. kindly provided the buprenorphine XRB (Ethiqa) for this study.

